# Novel Highly Homeostatic B Cells induced by Immunomodulatory Oligonucleotide IMT504

**DOI:** 10.64898/2025.12.18.694904

**Authors:** Juan Manuel Rodriguez, Fernanda Elias, María Victoria Oberholzer, Claudia Zylberberg, Juan Manuel Flo, Ricardo Agustín Lopez, Ana Julia Bridges, Diana Alicia Jerusalinsky, Jorge Zorzopulos

## Abstract

Cellular homeostasis depends on endoplasmic reticulum (ER) function and is highly sensitive to ER stress typically triggered by protein-folding overload. To restore balance, eukaryotic cells engage the unfolded protein response (UPR), a network of signaling pathways that enhances resistance to stress by upregulating chaperones, reducing global translation, and promoting degradation of unfolded proteins via autophagy. Additional stress-sensing pathways, including the NRF2 antioxidant response and the AMPK energy-sensing axis, converge to reinforce cellular homeostasis.

In pathological conditions, reinforcing immune homeostasis constitutes a promising therapeutic strategy. Among emerging pharmacological agents that activate endogenous protective pathways, the synthetic immunomodulatory oligonucleotide IMT504 has shown favorable pharmacokinetics and stability in preclinical models, although its mechanisms of action remain unclear.

Using purified human CD19⁺ B cells, we investigated the cellular, molecular, and transcriptional responses to IMT504 stimulation, to uncover new insights into its mode of action and broader implications for immune-mediated homeostasis.

We discovered that incubation of CD19⁺CD27⁻ B cells with IMT504 directly induces a-previously unreported-subpopulation of human B cells, which we termed **B homeostatic cells (Bhom)**. These Bhom cells display a distinctive phenotype (CD27⁻CD24⁺CD38⁺CD138⁻MUC1⁺) and upregulate genes associated with mitochondrial metabolism, proteostasis, antioxidant defense, and survival.

Promoter analysis of the most highly upregulated genes revealed an **Invariant Gene Transcriptional Regulator (IGTR)** signature with binding motifs for MYC, CREB1, CTCF, EP300 and NRF1/NRF2, suggesting a coordinated transcriptional control of the involved homeostatic programs. Network analysis linked NRF1 to SIRT1/PGC1α metabolic axis and NRF2 to IGTR-regulated factors. Furthermore, we found that IMT504 binds ATP-citrate lyase (ACLY), an endogenous repressor of AMPK, indicating that IMT504 may activate AMPK-dependent autophagy and downstream NRF1/NRF2 signaling.

Altogether, these findings identify Bhom cells as a novel B-cell subset that embodies homeostatic features and reveal IMT504 as a potent inducer of B-cell–mediated homeostasis through AMPK-driven transcriptional reprogramming. This mechanism strongly supports IMT504 as a promising candidate for immune-modulatory therapies aimed at restoring immune homeostasis.

**SIGNIFICANCE STATEMENT:** The discovery of a novel B-cell subset, termed *B homeostatic cells* (Bhom), reveals a previously unrecognized axis of immune response. Incubation of human CD19⁺CD27⁻B cells with the immunomodulatory oligonucleotide IMT504 directly induces **Bhom cells.** Its differentiation engages a conserved transcriptional network centered on AMPK–NRF1/NRF2 signaling, highlighting a potential therapeutic pathway to restore or enhance immune homeostasis.

## INTRODUCTION

Homeostasis, as defined by Walter Cannon (1871–1945), refers to the coordinated regulation of internal parameters—such as temperature, pH, and blood pressure—to preserve physiological equilibrium (1) According to Cannon, this constancy requires the activity of sensors capable of detecting and responding to deviations from set points. This concept remains widely accepted due to its simplicity and explanatory power.

At the cellular level, homeostasis refers to the balanced operation of molecular networks within functional limits. Mechanisms that ensure this balance include protein folding pathways assisted by chaperones, degradation of unfolded or misfolded proteins, autophagy, ATP-generating processes (especially those preserving mitochondrial integrity), antioxidant defenses and expression of cytoprotective proteins (2). Disruptions in these systems are associated with aging, trauma, and a broad range of diseases—including cancer, infections, metabolic syndromes, immune and neurological disorders (3).

Cellular homeostasis is closely linked to the endoplasmic reticulum (ER) and is highly sensitive to ER stress, typically caused by excessive protein-folding demands. In response, eukaryotic cells activate the unfolded protein response (UPR), a network of signaling pathways that enhances resistance to stress by upregulating chaperones, reducing global translation, and promoting degradation of unfolded proteins via autophagy (4, 5). In parallel, other stress sensors reinforce homeostasis through additional mechanisms: oxidative stress activates the NRF2 transcription factor which induces antioxidant responses (6), while energy depletion triggers AMPK, a kinase that restores ATP balance by modulating both anabolic and catabolic processes (7).

Under pathological conditions, homeostasis reinforcement is a desirable goal for therapeutic development. Two major strategies have been explored: transplantation of mesenchymal stem cells (MSCs), which secrete homeostasis-supporting factors (8), and pharmacological agents that activate endogenous protective pathways. Examples include the natural compounds curcumin (9) and flavonoids (10), as well as the synthetic drugs metformin (11) and the oligonucleotide (ODN) IMT504 (12). While preclinical studies using these compounds have shown success across various disease models, clinical translation has been limited by issues such as poor stability or bioavailability. IMT504 stands out for its synthetic stability and favorable pharmacokinetics (13). This belongs to the PyNTTTTGT ODNs class, that include at least one PyNTTTTGT octanucleotide in their sequence, and targets B-cells and/or plasmacytoid dendritic cells, leading to strong immunostimulation in humans and other primates (14). However, its mechanisms of action remain incompletely understood.

In this study, we investigated how IMT504 exerts its effects on immune B cell homeostasis. Using purified human CD19⁺ B cells, we examined the cellular, molecular, and transcriptional responses to IMT504 stimulation, aiming to uncover new insights into its mode of action and probable broader implications for immune-mediated homeostasis.

We showed that IMT504 directly promotes the emergence of a previously unreported subpopulation of human CD19⁺ B cells with a distinctive homeostatic phenotype, that we named ‘B homeostatic cells’ (Bhom).

## MATERIALS AND METHODS

### 1) Microarray Assays

#### B Lymphocyte Purification

Blood samples from healthy donors were obtained from the Hematology Division of Hospital Alemán (Buenos Aires, Argentina), using heparin as an anticoagulant. Peripheral blood mononuclear cells (PBMCs) were isolated by density gradient centrifugation with Ficoll-Hypaque (GE Healthcare Bio-Sciences, Uppsala, Sweden). Blood samples were diluted 1:1 in RPMI 1640 medium (Gibco, Thermo Fisher Scientific, Waltham, MA, USA) supplemented with 2 mM L-glutamine, 50 µg/ml gentamicin, and 25 mM HEPES, then centrifuged at 1,000 × g for 40 minutes at 20°C. The resulting PBMC pellet was washed and resuspended in RPMI 1640 medium supplemented with 10% fetal calf serum (FCS; Invitrogen, Thermo Fisher Scientific). CD19+ B lymphocytes were purified from PBMCs using CD19 MicroBeads (MACS, Miltenyi Biotec, Germany; Order No. 130-050-301). Cell purity exceeded 96%, as determined by flow cytometry.

#### Cell Treatment and Culture

Purified CD19+ B cells were cultured in 48-well plates at a density of 2 × 10⁶ cells/ml in 0.5 ml of RPMI 1640 medium per well. Cells were stimulated with IMT504 at a final concentration of 1.5 µg/ml. Control cells were incubated under identical conditions without IMT504. Samples were collected at 2, 4, and 22 hours post-treatment.

#### RNA Extraction and Microarray Processing

Total RNA was extracted from five independent CD19+ B cell cultures using the RNeasy Mini Kit (QIAGEN Inc., Germantown, MD, USA). RNA samples were pooled for downstream analyses. Gene expression profiling was conducted using CodeLink UniSet Human 20K Bioarrays (Amersham Biosciences, Buckinghamshire, UK), following the manufacturer’s protocols. RNA quality was confirmed via spectrophotometry and 1.2% agarose gel electrophoresis.

#### cRNA Preparation and Hybridization

First-strand cDNA was synthesized using 1 µg of total RNA with a T7 oligo(dT) primer at 42°C for 2 hours. Bacterial mRNAs were added as internal controls. Second-strand cDNA synthesis followed, and cDNA was purified using the QIAquick PCR Purification Kit (QIAGEN). In vitro transcription was performed at 37°C for 14 hours using biotin-11-UTP (PerkinElmer), and biotinylated cRNA was recovered with the RNeasy Mini Kit. Ten µg of cRNA were fragmented at 94°C for 20 minutes before hybridization.

#### Detection and Data Analysis

Hybridization was conducted at 37°C for 8 hours at 300 rpm. After post-hybridization washes, slides were incubated with Cy5-streptavidin, briefly rinsed in 2× SSC/0.05% TWEEN™, dried by centrifugation, and scanned using an arrayWoRx™ “e” scanner (Applied Precision, Issaquah, WA, USA). Data were analyzed with CodeLink Expression Analysis software v4.1. Only high-quality spots labeled “Good” were included. Differential expression was evaluated using Significance Analysis of Microarrays (SAM, Stanford University). Genes with a false discovery rate (FDR) of 0% and a >2.5-fold increase were considered significantly upregulated.

### 2) Cytological Assays

#### B Cell Isolation

For cytological and cytokine assays, B cells were purified from PBMCs using the B Cell Isolation Kit Human II (MACS, 130-091-151; Miltenyi Biotec). Purity was confirmed at 98% by flow cytometry.

#### Flow Cytometry

One × 10⁶ purified B cells were cultured in 96-well round-bottom plates (100 µl/well) at 37°C in 5% CO₂. Cells were stimulated with 6 µg/ml IMT504 or left untreated (controls). After incubation, cells were stained with fluorochrome-conjugated monoclonal antibodies (Miltenyi Biotec): CD24 PE-Vio615 (130-112-664; REA832), CD38 PE-Vio770 (130-108-838; REA572), and CD227 (MUC-1) APC (130-106-784; REA448). Staining was performed at 4°C for 10 minutes in the dark. Cells were washed, fixed in 1% formaldehyde, washed again, and analyzed on a FACScan II flow cytometer (Becton Dickinson). Viability was >85% after 72 hours. At least 5 × 10⁵ events/sample were analyzed using FlowJo v7.6. 3 independent assays were performed

### 3) Cytokine Secretion Assays

#### IL-10 and IL-35 Quantification

B cells (1 × 10⁶/well) were cultured as above. Supernatants were collected and analyzed using commercial ELISA kits: Human IL-10 ELISA Set (OptEIA™, BD Biosciences, Cat. No. 555157; detection limit: 31 pg/ml) and Human IL-35 ELISA Kit (Biomatik, Cat. No. EKC40941; detection limit: 62 pg/ml). 3 independent assays were performed.

### 4) Binding of PBMC Cytoplasmic Proteins to IMT504

#### Protein Capture and Isolation

Human or rat PBMCs were isolated via Ficoll-Hypaque density gradient centrifugation and incubated with 3′-biotinylated IMT504 for 1 hour at 37°C. Controls lacked IMT504. Cells (∼1 × 10⁷) were lysed in Brij-97 buffer with protease/phosphatase inhibitors. Lysates were centrifuged at 12,000 × g for 15 minutes at 4°C, and supernatants were subjected to streptavidin bead pull-down. Proteins were resolved by SDS-PAGE and stained. Differential bands were excised and processed for mass spectrometry.

#### Mass Spectrometry Analysis

Gel slices were discolored, reduced with DTT, alkylated with IAA, and digested with trypsin overnight. Peptides were extracted, dried, and stored at −80°C. Samples were analyzed using an Orbitrap Fusion Lumos mass spectrometer coupled to an Easy-nLC 1200 system. Separation was performed on a C18 column using a 5–35% acetonitrile gradient. Data were acquired and analyzed using MaxQuant v1.6.17.0 against the UniProt human database. Assays were performed twice.

### 5) Autophagy Assays

#### Cell Culture and Treatments

HeLa cells stably expressing EGFP-LC3 were cultured in MEM (Gibco, Cat. 12800-058) supplemented with 10% FCS (Gibco, Cat. 16000-044). To induce autophagy, cells were treated with 0.2 µM rapamycin for 2–4 hours. For IMT504 assays, cells were treated with 0.1 µM IMT504. Controls included rapamycin-only, Bafilomycin A1 (20 nM), and the unrelated oligonucleotide IMT022. Cells were fixed in 4% paraformaldehyde, washed, and mounted with ProLong™ Gold Antifade. Images were acquired on a Zeiss LSM 800 confocal microscope (1024 × 1024 pixels). EGFP-LC3 fluorescence was quantified using a custom Python v3.10 script.

Desalted phosphorothioate oligodeoxynucleotides (ODNs) were purchased from Operon Technologies (Alameda, CA). ODNs were suspended in depyrogenated water, tested for lipopolysaccharide (LPS) contamination using the Limulus amebocyte lysate (LAL) assay, and stored at 15 °C until use. Purity was evaluated by high-performance liquid chromatography (HPLC) and polyacrylamide gel electrophoresis (PAGE). Only preparations with a purity of ≥97% and undetectable LPS levels were used. All assays were performed in duplicate.

### 6) Statistical Analysis

Statistical analyses were performed using IBM® SPSS® Statistics software (IBM Corp., Armonk, NY, USA).

### 7) Additional Resources and Data Visualization Tools

#### Literature Databases

PubMed (https://pubmed.ncbi.nlm.nih.gov) and Google Scholar (https://scholar.google.com)

#### Charting Tool

DATAtab (https://datatab.net/statisticscalculator/charts/create-scatterplot)

#### Protein Interaction Networks

STRING database (https://string-db.org/) Gene/Protein Annotations: Harmonizome database (https://maayanlab.cloud/Harmonizome/)

## RESULTS

### IMT504 Induces Two Distinct Transcriptional Phases in Human CD19⁺ B Cells

To explore the immunomodulatory effects of IMT504, we performed genome-wide transcriptome analysis on purified human CD19⁺ B cells incubated either with or without IMT504 for 2, 4, or 22 hours. Transcripts showing a ≥2.5-fold induction and a moderate coefficient of variation at any time point were categorized according to their predominant biological functions (Table 1(A) and 1(B)).

**Table 1.**
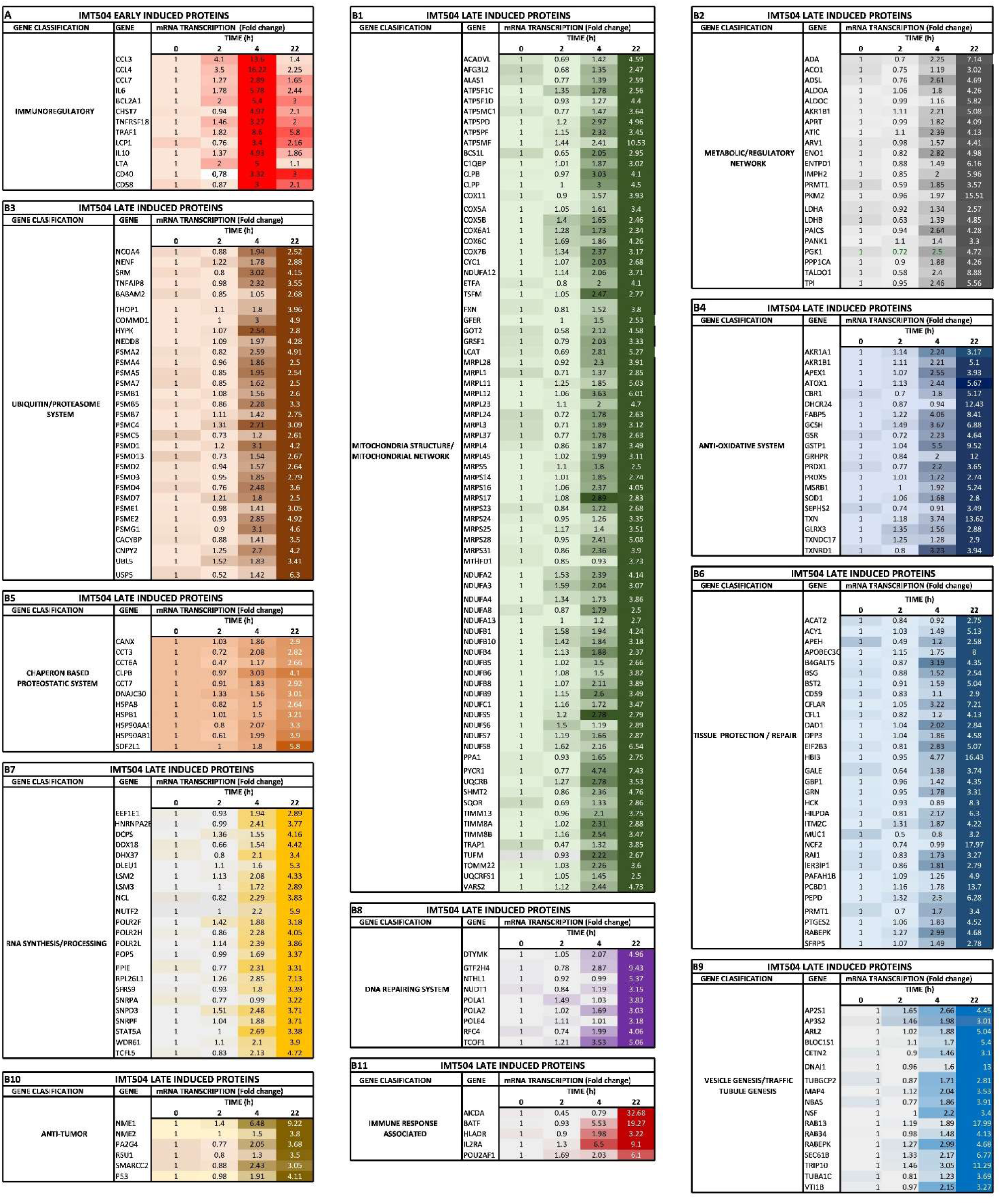
Time-dependent fold changes in gene expression in CD19⁺ B cells incubated with IMT504. Transcripts showing a ≥2.5-fold induction and coefficient of variation ≤30% at any time point were grouped according to shared biological functions or their involvement in specific cellular systems, based on classifications from scientific literature. Color intensity reflects the magnitude of fold change in mRNA expression at each time point (2, 4, and 22 hours) following IMT504 treatment of purified human CD19⁺ B cells.

During examination of the microarray data, it was observed that all induced genes could be firstly grouped according to two defined expression profiles that were named as early (Fig. 1(A)) and late (Fig. 1(B)) genes. Early transcripts, codify for a handful of mostly pro-inflammatory proteins (Table 1(A)), some of which have been previously shown to be induced by IMT504 acting on purified human B cells (14). On the other hand, late transcripts codify for a large number of proteins that can be grouped as components of sub-cellular structures and systems with strong homeostatic commitment, as presented in Table 1(B1-11). Coherently, cytokine secretion kinetics for IL-10 and IL-35—both anti-inflammatory mediators—paralleled the sequential expression of their corresponding genes (Figs. 1(C) and 1(D)).

**Figure 1.**
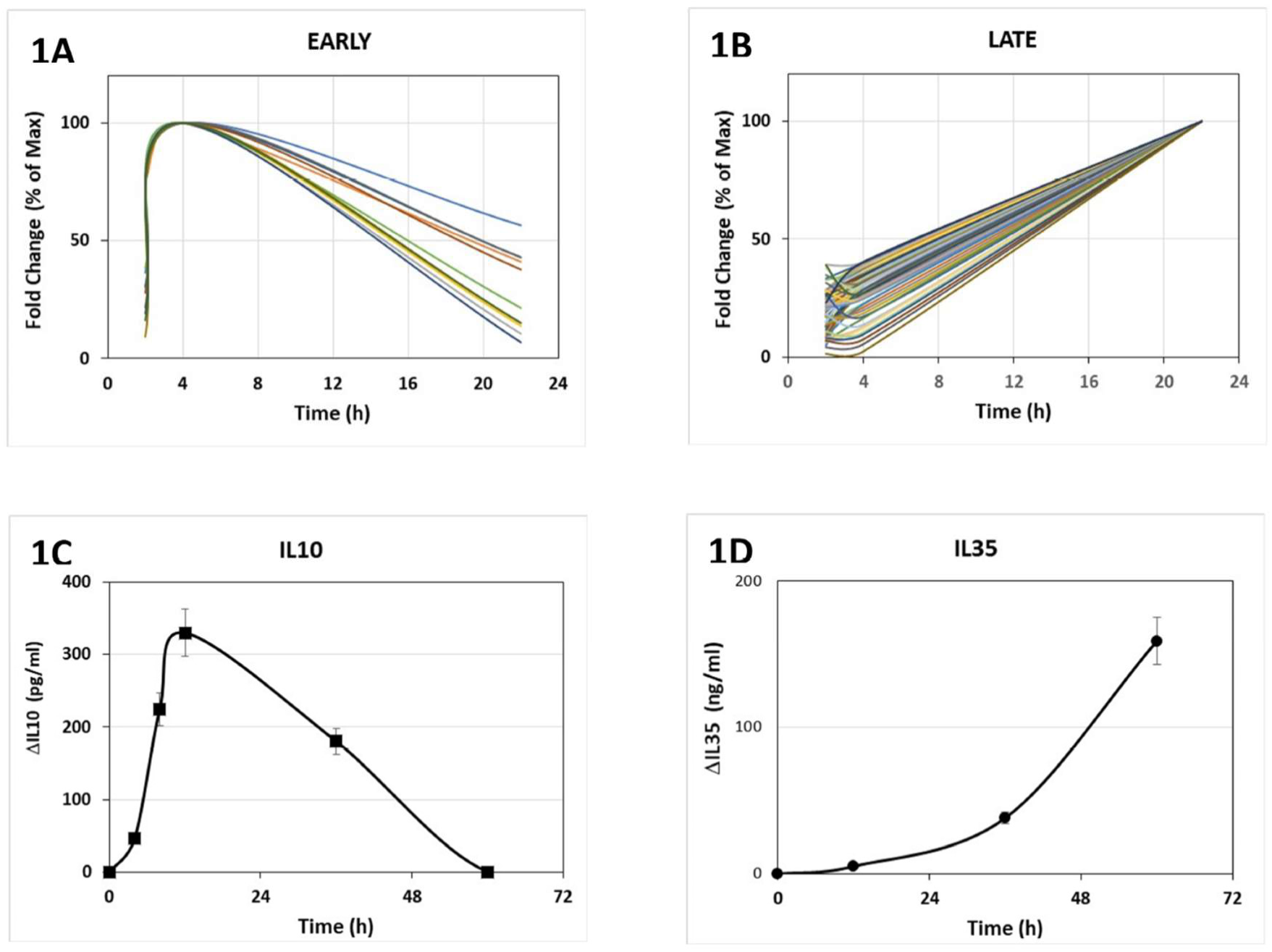
Gene expression and cytokine secretion in human CD19⁺ B cells stimulated with IMT504. (A) Early and (B) late mRNA expression patterns in CD19⁺ B cells incubated with IMT504, as defined in Figure 1. mRNA fold changes are expressed as a percentage of the maximum value observed for each gene across all time points. Gene expression in cells incubated without IMT504 showed no significant fluctuations and is not displayed. (C) IL-10 and (D) IL-35 secretion patterns measured in culture supernatants. Cytokine secretion is represented as the change (Δ) between successive time points.

Key pathways among the late-induced genes include the mitochondria (15, 16) (Table 1(B1)), the ubiquitin-proteasome system (17) (Table 1(B3)), molecular chaperones (18) (Table 1(B5)), and antioxidant defenses (19) (Table 1(B4)). Metabolic genes include those involved in purine synthesis (20) and glycolysis (21) (Table 1(B2)). Many late genes encode proteins involved in tissue protection and repair (14, 15) (Table 1(B6)). These findings indicate that after 22 h incubation with IMT504, a subset of the purified human blood CD19⁺B cells acquires a homeostatic transcriptional profile. This subpopulation was named “homeostatic B cells” or “Bhom” cells.

### Cytometric studies indicate that Bhom cells originate from CD19⁺CD27⁻ B cells. Phenotypic profile

To phenotypically characterize Bhom cells, flow cytometry was employed. MUC1, a surface protein whose transcript was significantly upregulated during late gene induction (Table 1(B6)), was selected as a Bhom marker candidate. Although MUC1 is typically linked to epithelial immune defense and inflammation regulation (22), its role in B cells remains unclear.

Freshly isolated CD19⁺ B cells displayed MUC1 positivity, which was almost lost after 12 h in culture in both, CONTROL conditions and IMT504 incubation (Fig. 2(I)). After 36 h of incubation, MUC1 expression continued to decline, though it was slightly increased after 84 h in CONTROL cultures. However, with IMT504 incubation a noticeable MUC1^+^ population emerges after 36 h of incubation, which is even greater after 84 h. These results strongly suggest that surface MUC1 immunoreactivity (MUC1⁺) represents a hallmark of Bhom differentiation, which is strongly induced by IMT504. It then also appears to be promoted by stress associated with prolonged culture. As it can be observed at the starting point MUC1^+^ B cells size is variable, though the small cells are predominant. Even after 12 h and 36 h incubation in control conditions, MUC1 marker mainly remains in small cells. And after 84 h of incubation, the slight increase in MUC1 expression in control cultures is predominantly located in small cells also. Accordingly, the populations expressing MUC1^+^ after 36 h and 84 h incubation in the presence of IMT504 also mainly consist of small cells.

**Figure 2.**
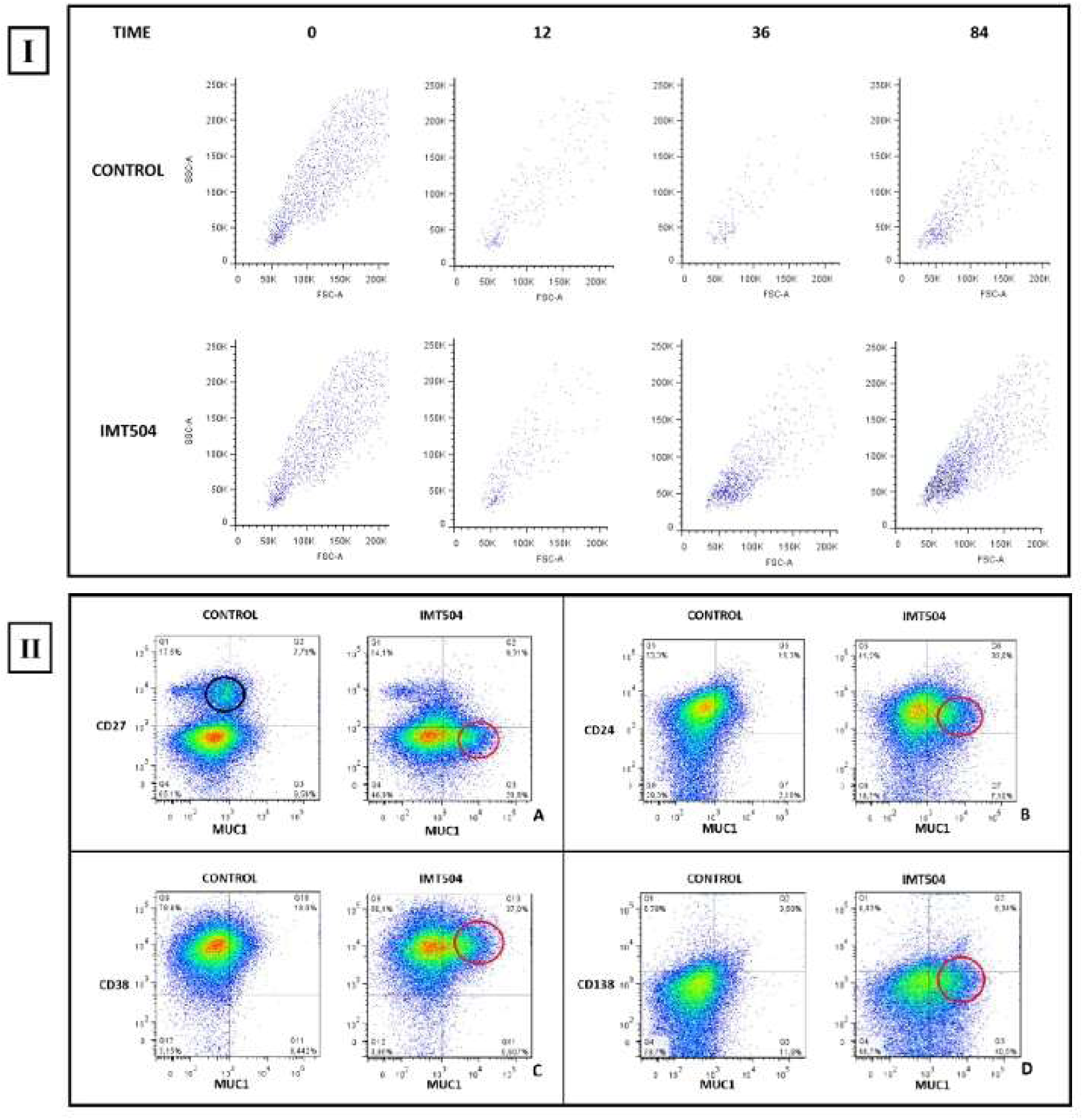
Cytometric analysis of CD19⁺ B cells following incubation with IMT504. I) Forward vs. side scatter (FSC vs. SSC) plots combined with MUC1 staining reveal the temporal dynamics of this marker expression in B cells incubated with or without IMT504 (CONTROL) for 12, 36, and 84 hours. After 36 hours of IMT504 exposure, a dense population of small MUC1⁺ cells become prominent, a population slightly detectable in control cultures. II) Flow cytometric analysis of MUC1 expression in combination with markers commonly used to define B cell subpopulations, after 36 hours of incubation with (IMT504) or without IMT504 (CONTROL): A) CD27 vs. MUC1: An abundant CD27⁻/MUC1⁺ subpopulation (red circle) appears after IMT504 treatment, accompanied by a small reduction in CD27⁺ cells (black circle). B) CD24 vs. MUC1: A prominent CD24⁺/MUC1⁺ subpopulation (red circle) emerges in IMT504-treated cells. C) CD38 vs. MUC1: An increased CD38⁺/MUC1⁺ subpopulation (red circle) is observed following IMT504 incubation. D) CD118 vs. MUC1: A distinct CD118⁻/MUC1⁺ subpopulation (red circle) becomes evident after treatment with IMT504.

We used well-known B cell markers —CD24 and CD38 (commonly used to define B cell developmental stages) (23), CD27 (a memory B cell marker) (24), and CD138 (a plasma cell marker) (25)— alongside MUC1, to further define Bhom phenotype. At the starting point, cell cultures displayed a similar distribution of these cell markers as that of the 36 h incubation control cultures (data not shown). A distinct CD27⁻MUC1⁺ population was identified after 36 h of IMT504 incubation (Fig. 2(II)A, IMT504, red circle), almost absent in untreated controls (Fig. 2(II)A, CONTROL). The fact that the difference between CD27⁻MUC1⁺ cells in both control and IMT504 incubated conditions (Q3IMT504-Q3CONTROL ∼20.2%, Fig. 2(II)A), and the difference between CD27⁻MUC1⁻ cells in both conditions ([Q4CONTROL-Q4IMT504 ∼18.2%], Fig. 2(II)A) appear to be similar, suggests lineage derivation. A reduction in CD27⁺ cells was also noted (Fig. 2(II)A, black circle), which will be discussed further on.

Additionally, subpopulations can be recognized in CD24⁺MUC1⁺ ([Q6IMT504-Q6CONTROL∼18 %], Fig. 2(II)B, red circle), CD38⁺MUC1⁺ ([Q10IMT504-Q10CONTROL∼19 %], Fig. 2(II)C, red circle), and CD138⁻MUC1⁺ ([Q3IMT504-Q3CONTROL∼29 %], Fig. 2(II)D, red circle) gates. It must be noticed that CD138⁺MUC1⁺ population almost did not change between IMT504 and control conditions ([Q1+Q2IMT504∼10,77 %; Q1+2CONTROL∼10,28 %], Fig. 2(II)D), excluding a direct link between Bhom and plasma cells.

These observations indicate that Bhom cells can be characterized by CD27⁻, CD24⁺, CD38⁺, CD138⁻, and MUC1⁺ markers after 36 h incubation. The co-expression of CD24 and CD38 at high levels on the precursor population is consistent with a transitional B-cell origin, suggesting that Bhom cells act as a novel differentiation fate for this developmentally poised subset. Approximately, 18-19 % of CD19⁺ cells differentiated into MUC1⁺ Bhom after 36 h of IMT504 exposure. However, for CD138 vs MUC1 marker combination a larger % was observed.

### IMT504 modifies additional CD19⁺ B cell subsets beyond Bhom differentiation

To further elucidate the effects of incubation with IMT504 on CD27, CD38 and CD59, CD19^+^ B cell expression, firstly, cell division by carboxyfluorescein-succinimidyl ester (CSFE) dilution as a function of CD27 expression was analyzed (after 96 h incubation, with IMT504 or without - CONTROL-) (Fig. 3(A)). Red arrows indicate the most noticeable cell populations that suffered division: around 1.97 % of cells in control conditions, and around 7.41 % when incubated with IMT504. Therefore, only ∼6% of CD19⁺ cells entered cell division by 96 h of incubation with IMT504.

**Figure 3.**
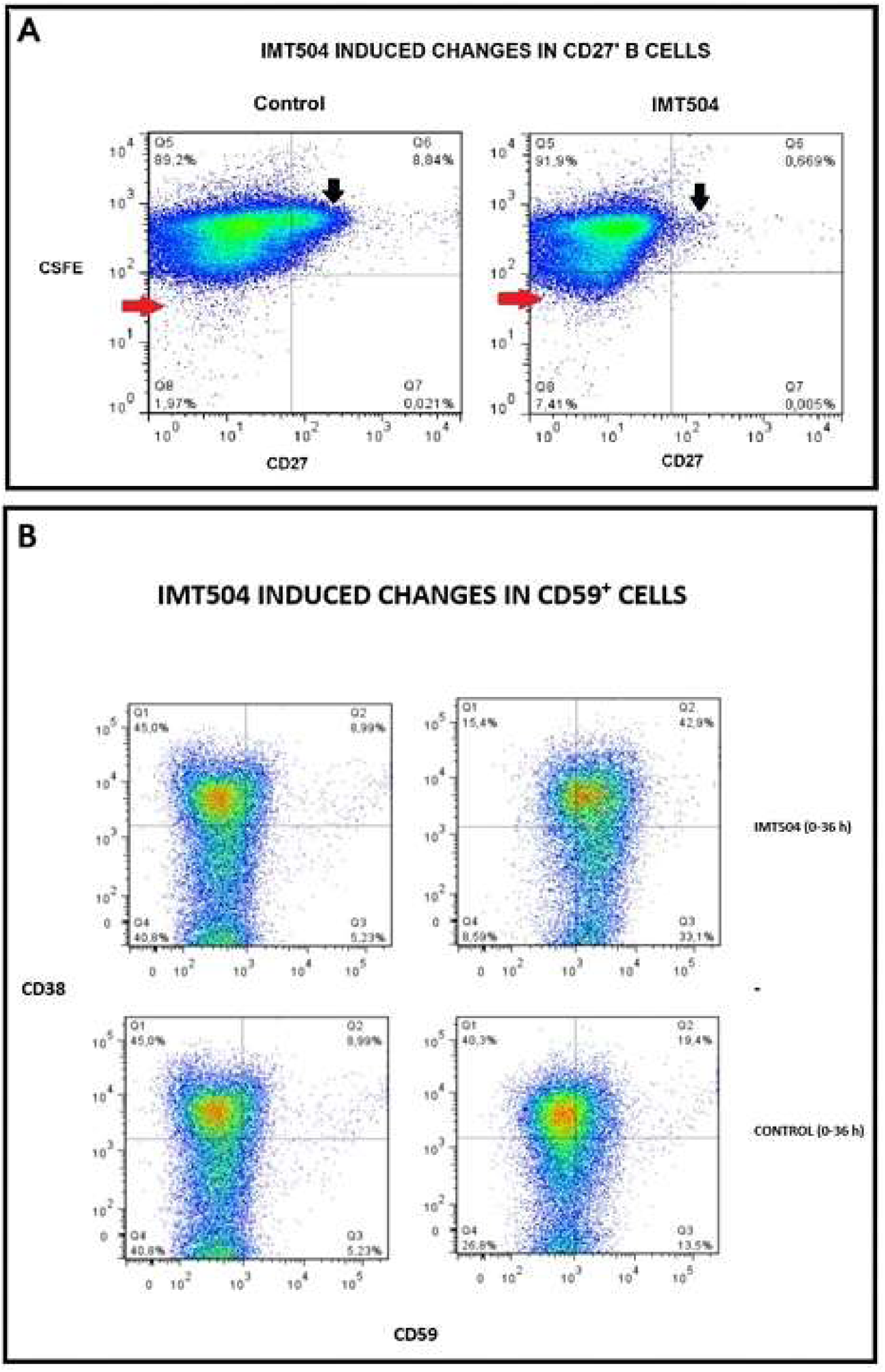
Effects of IMT504 on CD19⁺ B cell proliferation and expression of CD27 and CD59 markers. A) CD19⁺ B cell proliferation was assessed after 96 hours of incubation with or without IMT504 using carboxyfluorescein succinimidyl ester (CFSE) dilution in combination with CD27 staining. The red arrow highlights a population of proliferating cells in IMT504 treated cells and its absence in control conditions. The black arrow indicates a CD27⁺ subpopulation in control cells and its absence in IMT504 treated cells. B) CD38 expression as a function of CD59 expression, before (0 h) and after 36 h incubation with or without IMT504 (CONTROL).

Black arrows indicate CD27^+^ cell populations, which were about 8.84 % of cells in control conditions; a similar proportion was observed at the starting point (data not shown). CD27^+^ marker was around 0.669 % after incubation with IMT504 (Fig. 3(A)). As a result, approximately 8 % fraction of CD27⁺ cells lost its expression, suggesting either down-modulation of CD27 or dilution of the marker by specific proliferation of memory B cells during the initial proinflammatory phase (Table1(A)).

We next examined the effects of IMT504 incubation on CD59 expression as a function of CD38 expression. Approximately 19 % of CD19⁺ B cells became CD59⁺ after 36 h incubation in control conditions, while ∼ 62 % of CD19⁺ B cells became CD59^+^ after 36 h with IMT504 (CD59^+^ cells [Q2+Q3]: ∼ 14 % at the starting point, ∼76 % with IMT504 and ∼33 % in control conditions after 36 h incubation, Fig. 3(B)). Meanwhile, CD38 expression was not substantially modified (CD38^+^ cells [Q1+Q2]: ∼ 54 % at the starting point, ∼ 58 % after IMT504 incubation and ∼ 60 % after 36 h in control conditions; Fig. 3(B)).

This experiment reveals that incubation with IMT504 substantially stimulates CD59 expression in more cells (∼ 62 %) than only Bhom cells (∼ 18-29 %), indicating a broad response that extends beyond them. It is worth noticing that the incubation in control conditions was able to induce CD59 expression though in a smaller subpopulation.

### An invariant set of shared transcription regulators (IGTR) coordinate gene expression during CD19^+^B cell differentiation into Bhom cells

To investigate the genetic mechanisms underlying the differentiation of human CD19⁺ B cells into Bhom cells, we analyzed transcriptional regulatory elements using the ENCODE encyclopedia, focusing on promoter regions of genes upregulated by IMT504. This analysis revealed a consistent group of transcriptional regulators, hereafter referred to as the invariant gene transcriptional regulators (IGTR), comprising the following elements: **CHD1, CHD2, CREB1, CTCF, EP300, H2AFZ, JUND, MAX, MXI1, MYC, POLR2A, RAD21, SIN3A, TAF1, TBP, YY1**, and **ZNF143.**

Among these, **CREB1, JUND, MYC, TBP, TAF1**, and **YY1** are well-established transcription factors, with **YY1** also functioning as an epigenetic regulator (26). **H2AFZ** is a histone variant with key roles in mitochondrial gene transcription (27), while **POLR2A** encodes the largest subunit of RNA polymerase II. Other members, including **CHD1, CHD2, CTCF, RAD21, SIN3A**, and **ZNF143** are epigenetic regulators that modulate transcription primarily through changes in chromatin topology (28, 29, 30, 31, 32). **EP300** encodes the histone acetyltransferase p300, a key chromatin remodeler (33), and **MAX** and **MXI1** are known MYC co-regulators (34).

Figure 5(A) presents a STRING network of the IGTR components, illustrating their potential physical and/or functional interconnections during CD19⁺ B cells to Bhom cell differentiation. The tight connectivity among these regulators suggests a coordinated transcriptional program.

To further explore the transcriptional patterns, violin plots (Fig. 4(B)) and distribution analyses (Figs. 4(C–D)) of late gene fold changes allow to identify two expression patterns, each comprising three functional gene classes: “red” (CHAP, MIT, PROT) and “blue” (ANTI-OX, MET, TP) clusters. An independent t-test comparing red and blue patterns showed a significant difference in mean fold changes (t(193) = –5.88, p < 0.001), with the blue group showing greater upregulation and higher variability (For further details on Fig.4(B) statistics see Supp. Fig S1).

**Figure 4.**
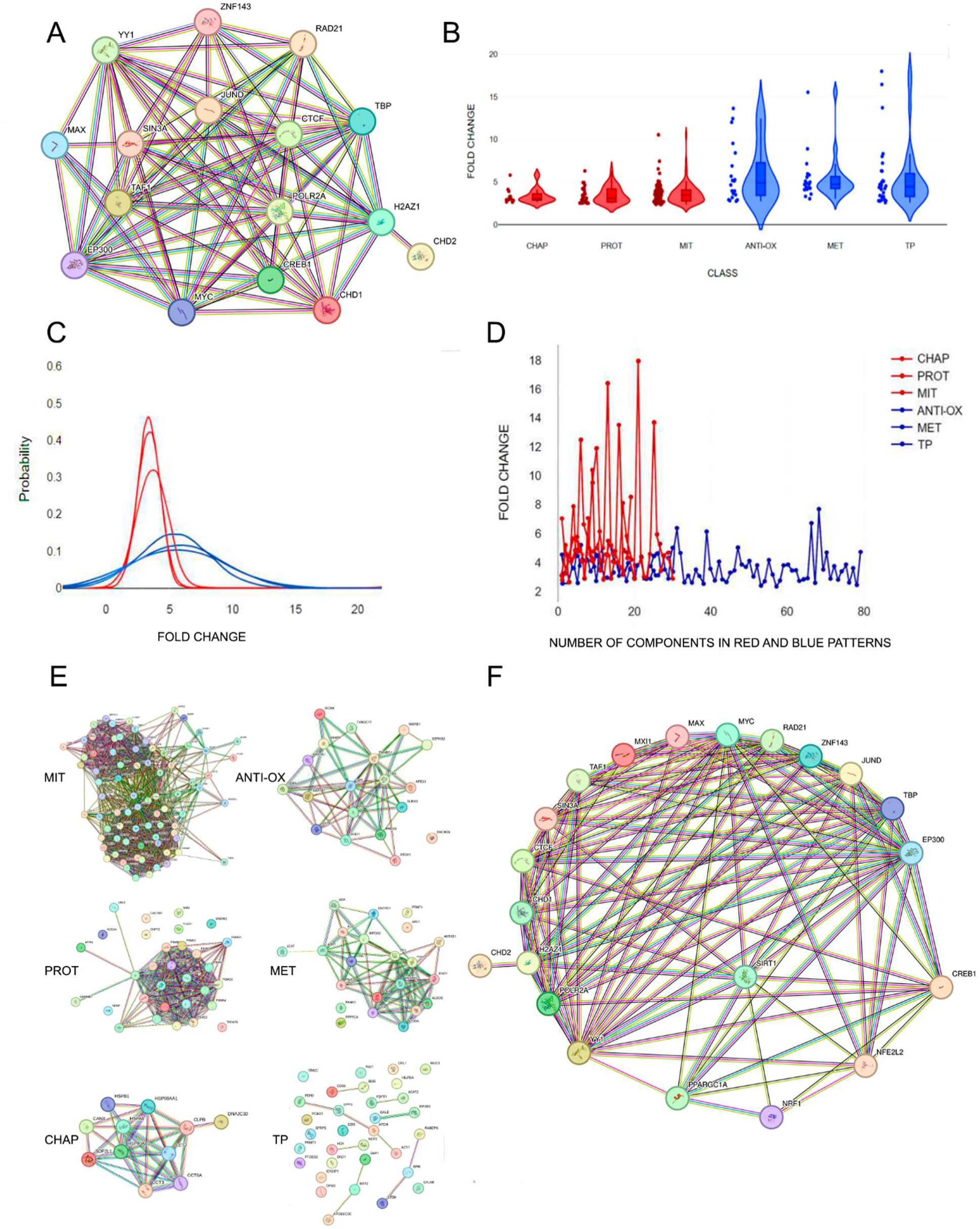
Analysis of the Invariant gene transcriptional regulatory (IGTR) elements in the promoters of genes induced during Bhom cell differentiation, and their relationship to known transcriptional regulators of cellular homeostasis. A) STRING interaction network of IGTR components, highlighting their connectivity. B) Violin plots of fold changes in late-transcript genes from the main homeostatic protein groups-as listed in Figure 1-, revealing two distinct expression patterns (red and blue), each comprising three gene categories: Chaperones (CHAP), Proteasome (PROT), Mitochondrial (MIT) and Antioxidant (ANTI-OX), Metabolic (MET), and Tissue Protection (TP), respectively. C) Normal distribution plots confirming the segregation of genes into the two distinct expression patterns. D) Line plots of fold-change dynamics further supporting the classification into red and blue gene expression clusters (see Results section). E) Intra-group network analysis of genes within the red and blue clusters, illustrating internal connectivity and functional cohesion. F) Network diagram depicting functional links between IGTR components and key transcriptional regulators involved in cellular homeostasis: NRF1, NFE2L2 (NRF2), and the co-activators SIRT1 and PGC1α.

One-way ANOVAs found no significant intragroup differences in either the red (F(2,119) = 0.65, p = .524) or blue (F(2,70) = 0.16, p = .857) clusters, supporting pattern coherence. The **red cluster** includes genes encoding structural components of major subcellular complexes (e.g., mitochondrial components, proteasomes), suggesting tight stoichiometric constraints on their expression. STRING analysis revealed strong connectivity among these genes, consistent with coordinated synthesis. In contrast, the **blue cluster** encompasses genes related to antioxidant defense, metabolism, and cytoprotection. These functions typically require more flexible regulation and showed weaker STRING connectivity, suggesting looser coordination during stress responses (Fig. 4(E)). STRING cluster analysis of all genes expressed after 22 hours of IMT504 exposure strongly supports this distinction (Supplemental Material Fig. S2).

With regards to the identity of potential master regulators of each cluster, literature indicates that **NRF1** most likely governs the red cluster (35, 36, 37), while **NRF2** is the main regulator of the blue cluster (38, 39, 40). This led us to examine whether these factors interact with IGTR components.

We generated a new STRING network (Fig. 4(F)) that included IGTR elements, **NFE2L2 (NRF2), NRF1**, and their common co-activators **SIRT1** and **PPARGC1A (PGC1α)**(41, 42). This analysis revealed:

a) **SIRT1** interacts with several IGTR components and with **PGC1α**,
b) **NRF1**shows limited interaction with IGTR,
c) **NRF2** significantly interacts with **JUND, TBP, YY1, MYC**, and **EP300**,
d) **NRF1**is functionally linked to **PGC1α**, and
e) **PGC1α** also connects with **EP300** and **CREB1**.

Together, these findings suggest that **NRF2** operates within a tightly regulated network of transcriptional co-regulators that may fine-tune stress-specific responses, while **NRF1** is also influenced-though indirectly-via the **SIRT1/PGC1α** axis. This transcriptional architecture may underlie the distinct regulatory logic governing homeostatic gene expression during IMT504-induced B cell differentiation. It is worth noting, that in a random sample of 200 genes retrieved from the ENCODE encyclopedia, only 27.5% simultaneously contained both the IGTR and the NRF1/2 binding sequence in their promoter regions. In contrast, this fraction amounted 94% in an equal sized sample of genes upregulated during Bhom differentiation induced by IMT504. A binomial test confirmed that this enrichment is highly significant (p ≈ 5.2 × 10⁻⁸⁹).

### IMT504 binds ACLY relieving its AMPK inhibition likely intiating Bhom differentiation

Our working hypothesis proposes that, to initiate Bhom differentiation, a master regulator upstream of SIRT1, PGC1α, NRF1, and NRF2 should be activated —either directly or indirectly— by IMT504. To explore how IMT504 started to induce such differentiation of CD19⁺ B cells into Bhom, human B cells were incubated with biotin-conjugated IMT504, lysed, and subjected to membrane solubilization using Brij97 detergent, followed by centrifugation. IMT504-bound proteins were then isolated from the resulting supernatants using streptavidin-Sepharose beads. The precipitated proteins were separated by gel electrophoresis; prominent bands, absent in control samples, were excised for identification by mass spectrometry (Fig. 5(I)). Most IMT504-bound proteins were identified as common nucleic acid-binding proteins unrelated to SIRT1, PGC1a, NRF1, or NRF2. However, one of IMT-504-bound proteins stood out: ATP-citrate lyase (ACLY), an enzyme that plays a crucial role in the synthesis of acetyl-CoA and which is also known as an AMPK inhibitor (43) (Fig. 5 (I)A).This strongly suggests that IMT504 binding to ACLY suppressess its AMPK-inhibitory activity, thereby allowing AMPK activation, a potential key step in initiating CD19^+^ B cell differentiation into Bhom.

**Figure 5:**
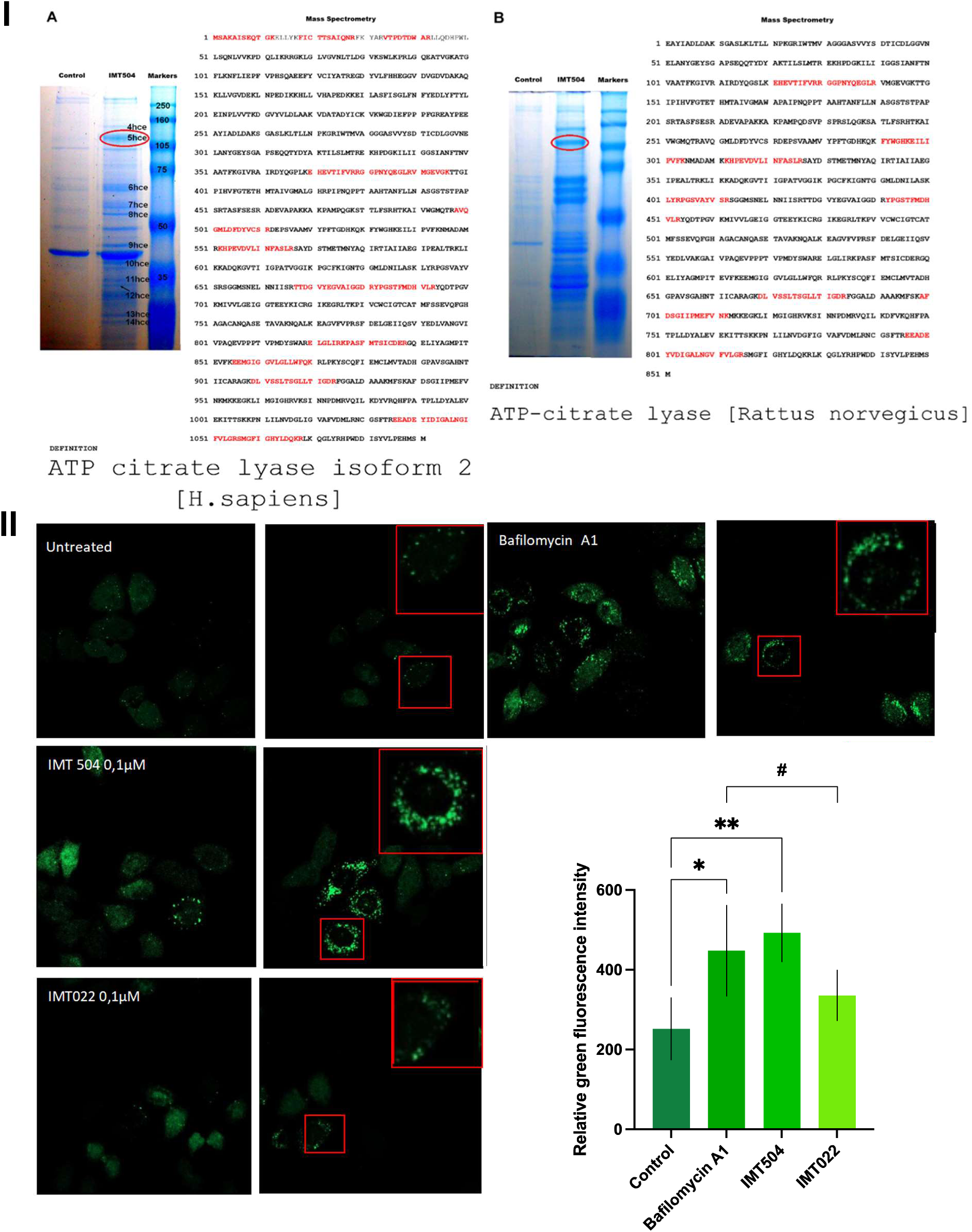
IMT504 binds to ACLY and stimulates autophagy. I) Binding of IMT504 to ACLY in B cell cytoplasmic extracts. Gel electrophoresis of cytoplasmic protein extracts from A) human and B) rat CD19⁺ B cells incubated with biotin-labeled IMT504 or left untreated (Control). The red circle is pointing out ATP-citrate lyase (ACLY) band, as identified by mass spectrometry. This band is absent in control samples, indicating specific binding of IMT504 to ACLY. II) Autophagy activation by IMT504. Representative fields of cultured cells in each condition. Red square insets: higher-magnification view of representative cells. Bottom right: total fluorescence quantification graph. One way ANOVA, Dunnett’s multiple comparisons test, * p < 0.05, ** p< 0.01. Unpaired t test, # p < 0.05.

Since it is well-known that AMPK activation stimulates autophagy (44, 45), we further investigated whether IMT504 could induce it. HeLa cells expressing EGFP-LC3 were incubated with either IMT504, IMT022 (same length ODN as IMT504, but with a different sequence), rapamycin, a partial activator of baseline autophagy, or Bafilomycin A1 as a positive control (inhibitor of autophagosome-lysosome fusion that leads to vesicle accumulation). Fluorescent labelling showed that IMT504 induced strong autophagosome accumulation, similar to that of Bafilomycin A1, surpassing that of control with rapamycin. Fluorescence was also higher than that following IMT022 incubation (Fig. 5(II)). Therefore, IMT504 strongly induced autophagic activity in HeLa cells. These data support a model in which IMT504 binds to ACLY relieving its AMPK inhibition, thereafter initiating autophagy and potentially triggering differentiation of a subpopulation of CD19^+^cells into Bhom.

## DISCUSSION

Several years ago, our group reported that IMT504, an immunomodulatory ODN lacking CpG motifs, could stimulate the expansion of mesenchymal stem cell (MSC) precursors both *in vitro* and *in vivo* (46). Independently, numerous studies have demonstrated that MSC transplantation effectively treats a wide range of pathological conditions in animal models of human disease (47). These observations led us to hypothesize that IMT504 treatment might itself confer therapeutic benefits in such models by promoting endogenous MSC activity. Supporting this hypothesis, independent research teams have shown that IMT504 treatment yields beneficial effects in diverse pathological conditions, including diabetes (46), neuropathic pain (47), chronic inflammation (48), sepsis (49), multiple sclerosis (50), and liver fibrosis (51). These findings suggested that the therapeutic effects of IMT504 might be primarily mediated through the promotion of endogenous MSC expansion (12).

The present study reveals a previously unrecognized mechanism of action of IMT504 through direct induction of a novel subpopulation of human CD19⁺B cells with a strong homeostatic profile, which we named “B homeostatic cells” or just Bhom.

Transcriptomic profiling after (22 h) incubation with IMT504, further revealed that Bhom cells express a wide array of genes associated with cellular homeostasis, including antioxidant proteins, chaperones, and mitochondrial regulators (Table 1). After incubation with IMT504, two temporarily discriminated gene expression patterns appeared: early (Fig. 1(A)) and late (Fig. 1(B)). Early transcripts codify for a handful of immunomodulatory proteins; some have already been reported as induced by IMT504 acting on purified human B cells (14). On the other hand, late transcripts codify for a large number of proteins that are components of sub-cellular structures and systems with relevant homeostatic functions (Table 1). Late-induced genes include several key pathways consisting of those controlling selective groups of genes during cellular stress (15), or the generation of ROS mediated signals regulating immune responses and autophagy in the mitochondria (16) (Table 1(B1)); the ubiquitin-proteasome system (17) (Table 1(B3)), molecular chaperones (18) (Fig. 1(B5)), and antioxidant defenses (19) (Table 1(B4)). Also, some metabolic genes are included (i.e., those involved in purine synthesis and glycolysis, associated with purinergic signaling (20) and lactate shuttling (21)) (Table 1(B2)). Moreover, some late genes encoded proteins involved in tissue protection and repair (14, 15) (Table 1(B6)). These findings indicate that, after the incubation with IMT504, a subset of purified human CD19⁺B cells acquires the homeostatic transcriptional profile leading to “Bhom”.

Flow cytometric analysis revealed that Bhom cells differentiate from transitional B-cell precursors, acquiring a distinct CD27⁻CD24⁺CD38⁺CD138⁻MUC1⁺ phenotype after 36 hours of incubation with IMT504 (Fig. 2 (I) and (II)). A key defining event was the strong upregulation of MUC1, a molecule implicated in tissue protection, establishing it as a hallmark of Bhom differentiation (52). The absence of the plasma cell marker CD138 definitively distinguished Bhom cells from terminal antibody-secreting lineages. Collectively, this unique surface profile, combined with functional characteristics such as higher IL-35 expression (Fig. 1(D)), establishes Bhom cells as a novel immunoregulatory population. The broad response observed in changes of CD59 expression after incubation with IMT504 indicates that it potentially triggers the activation of physiological defensive mechanisms beyond the generation of Bhom cells (Fig. 3). Similar to MUC1 (Fig. 2(I)), a mild though noticeable increase in CD59 expression was also seen in control cultures (Fig. 3(B)). This suggests that the cellular stress involved in the culturing process itself would trigger changes in B cells resembling those leading to Bhom.

Microarray analysis revealed that Bhom differentiation is characterized by the coordinated transcription of several genes encoding proteins with well-established homeostatic functions. These proteins fall into multiple functional categories, including antioxidants, molecular chaperones, and mitochondrial components. Because this transcriptional response appeared to be coordinated by a unique upstream signal, we searched for common regulatory elements in the promoter regions of Bhom-associated genes. There were transcription factors such as CREB1, MYC, JUND, YY1, TBP, TAF1, epigenetic regulators like CDH1, CDH2, CTCF, RAD21, SIN3A, ZNF143, the chromatin modifier EP300, and core machinery components such as H2AFZ and POLR2A. From the STRING analysis emerges that these genes form a tightly interconnected regulatory network, which might function as a coordinated complex (Fig. 4(A)). For instance, MYC-known to participate in diverse transcriptional programs (53)-might be a central node within this network (Fig.4(A) and (F)).

From violin plots (Fig. 4(B)) and distribution analyses (Figs. 4(C–D)) of late gene expression fold changes, two expression patterns were identified: “red” (CHAP, MIT, PROT) and “blue” (ANTI-OX, MET, TP). These patterns showed a significant difference in mean fold changes, with the blue group showing greater upregulation and variability (See Supplementary Table S1 for statistical analysis). Red-pattern genes encode mitochondrial components, proteasomes and chaperones, which are structurally related homeostatic proteins, suggesting a tight related regulation. Blue-pattern genes encoded proteins involved in metabolic or cytoprotective functions, with looser regulatory constraints (Fig. 4(E)).

Based on their expression patterns, and according to literature, these genes likely correspond to gene programs regulated by NRF1 and NRF2. If so, this would imply that the IGTR network communicates functionally with these transcription factors to regulate late-stage gene expression during Bhom differentiation. In the putative regulatory crosstalk revealed by STRING analysis between the IGTRs and the transcription factors involved, SIRT1 – which has been reported to mainly enhance NRF2 activity (54, 55, 56)-appears to bridge components of IGTR with NRF1/2 pathways. NRF1 interacts mainly through SIRT1, while NRF2 also interacts with several IGTR members (i.e., MYC, EP300, JUND, YY1, and TBP), in a complex and context-dependent way (Fig.4(F)). MYC, for instance, can both repress and enhance NRF2-mediated transcription (57, 58), while EP300 promotes this transcription-via chromatin remodeling and transcriptional co-activation-(59, 60, 61). The diverse interaction shown by the STRING analysis might enable flexible cell-specific stress responses. This interpretation aligns with our findings and prior work showing that IMT504 exerts distinct effects in different cell types (14, 62, 63, 64).

Interestingly, all late-expressed Bhom related genes contain TBP binding sites, suggesting TATA-box-mediated regulation, typically linked to stress-responsive genes activation (65). This observation underscores the adaptive transcriptional flexibility of Bhom cells in response to environmental cues.

Following our working hypothesis, -that to initiate Bhom differentiation, a master regulator upstream of SIRT1, PGC1α, NRF1, and NRF2 should be activated by IMT504-, CD19⁺ B cells were incubated with the ODN, and proteins bound to it were isolated. Most of the proteins identified were common nucleic acid-binding proteins unrelated to the SIRT1–PGC1α–NRF1/2 axis. However, ATP-citrate lyase (ACLY), a multifunctional enzyme also known to act as an AMPK inhibitor (43), surged as a possible relevant step in IMT504 mechanism (Fig. 5 (I)). Concerning IMT504’s effects on functional mechanisms, it induced strong autophagosome accumulation surpassing that of rapamycin (Fig. 5 (II)). Therefore, IMT504 binding to ACLY would suppress its AMPK-inhibitory activity, supporting a model in which IMT504 impedes or decreases AMPK inhibition, leading to initiate autophagy and Bhom differentiation.

Additional steps such as AMP binding or AMPK phosphorylation might also be required for full activation and warrant investigation.

This likely represents a key step in starting the differentiation of CD19⁺ B cells into Bhom. Further assays would be necessary to confirm this key role of AMPK.

Therefore, mechanistically our data point to AMPK activation as a key upstream driver of Bhom differentiation. Proteins binding to IMT504 included ATP-citrate lyase (ACLY), a multifunctional enzyme known to inhibit AMPK (43). We propose that IMT504 binding to ACLY releaves this inhibition, thereby facilitating AMPK activation. Consistent with this, IMT504 strongly induced autophagosome accumulation, surpassing the effect of rapamycin. Together, these findings support a model in which IMT504 releaves ACLY-mediated AMPK inhibition, initiating autophagy and activating NRF1/2-driven transcriptional programs that establish the Bhom phenotype. Further assays will be required to confirm AMPK’s central role, but this model provides a unifying framework linking IMT504 activity with Bhom differentiation.

Importantly, AMPK activation is known to be beneficial in many pathological contexts, including sepsis (66), diabetes (67), chronic pain (68), multiple sclerosis (69), and osteoporosis (70). Thus, AMPK-mediated induction of Bhom cells may explain IMT504’s broad therapeutic efficacy. This mechanism also aligns with IMT504’s pleiotropic effects in other cell types, including plasmacytoid dendritic cells (14), NK/NKT cells (62), MSCs (63), and pancreatic β cells (64).

### Projection

The robust homeostatic phenotype of Bhom cells suggests they are well-equipped to modulate tissue environments under stress. For this, intercellular communication via secreted factors, membrane-bound proteins, extracellular vesicles, or tunneling nanotubes may be critical (50). Whether Bhom cells employ such mechanisms remains unknown and is a key avenue for future investigation.

Exploring Bhom’s longevity, expansion *in vivo* and therapeutic potential when transplanted in isogenic models, would be of main importance. The observed benefits in a multiple sclerosis study (50) point to isogenic Bhom cell transplantation as a promising therapeutic strategy that deserves continued investigation.

Finally, it will be essential to determine whether Bhom cells are induced in vivo under physiological or pathological conditions, and to clarify how they interact with other homeostatic populations such as MSCs, Tregs, and M2 macrophages.

The favorable safety profile and scalability of IMT504 strongly support its continued development—either as a therapeutic agent capable of directly inducing Bhom cells in vivo, or as a platform for generating Bhom-based cell therapies.

## CONCLUSION

The discovery of specific B cells bearing a robust homeostatic profile defines a new cellular category named Bhom, and introduces new possibilities for autologous or allogeneic cell therapies (e.i. as it is suggested by preclinical data from a rat model of multiple sclerosis (50)).

The immunomodulatory IMT504 effectiveness across diverse disease models could be explained through the induction of Bhom cells and the activation of a homeostatic cellular network capable of responding to stressful conditions. The proposed mechanism of action sheds light onto the development of new therapeutic strategies for diseases that currently lack effective treatments, particularly those driven by chronic inflammation or impaired tissue resilience.

## Acknowledgments

We gratefully acknowledge Dr. Mario Rossi (Translational Medicine Research Institute—IIMT, CONICET, Facultad de Ciencias Biomédicas, Universidad Austral, Buenos Aires, Argentina) for generously providing the EGFP-LC3 HeLa cell line. We thank Ramiro Aníbal Carreño for his valuable assistance in figures design. This study was funded by Immunotech SA and Pablo Cassará SRL, Buenos Aires, Argentina. J.Z. and R.A.L. are shareholders of Immunotech SA. The remaining authors declare no conflicts of interest.

