## Supplemental material for "Novel Highly Homeostatic B Cells induced by Immunomodulatory Oligonucleotide IMT504"

Table 1. Comparison of Fold Change Between Red and Blue Patterns (Independent Samples t-Test).

| Group | Mean Fold Change (M) | SD | Variance | t (df = 193) | p-value | Levene's F | Levene's p-value |
|---|---|---|---|---|---|---|---|
| Red | 3.60 | 1.17 | 12.96 | -5.88 | < 0.001 | 38.83 | < 0.001 |
| Blue | 5.57 | 3.39 | 31.02 | | | | |

Table 2. ANOVA and Variance Test for Red Pattern Classes

| Class | Mean Fold Change (M) | SD | ANOVA F (2,119) | p-value | Levene's Test (Homogeneity) |
|---|---|---|---|---|---|
| CHAP | 3.39 | 0.93 | 0.65 | 0.524 | No significant difference |
| MIT | 3.69 | 1.27 | | | |
| PROT | 3.45 | 0.97 | | | |

Table 3. ANOVA and Variance Test for Blue Pattern Classes

| Class | Mean Fold Change (M) | SD | ANOVA F (2,70) | p-value | Levene's Test (Homogeneity) |
|---|---|---|---|---|---|
| ANTI-OX | 5.91 | 3.46 | 0.16 | 0.857 | No significant difference |
| MET | 5.33 | 2.67 | | | |
| TP | 5.52 | 3.86 | | | |

**Figure S1: Statistical analysis on the relationship of gene patterns and classes shown in Figures 5B, 5C and 5D).**

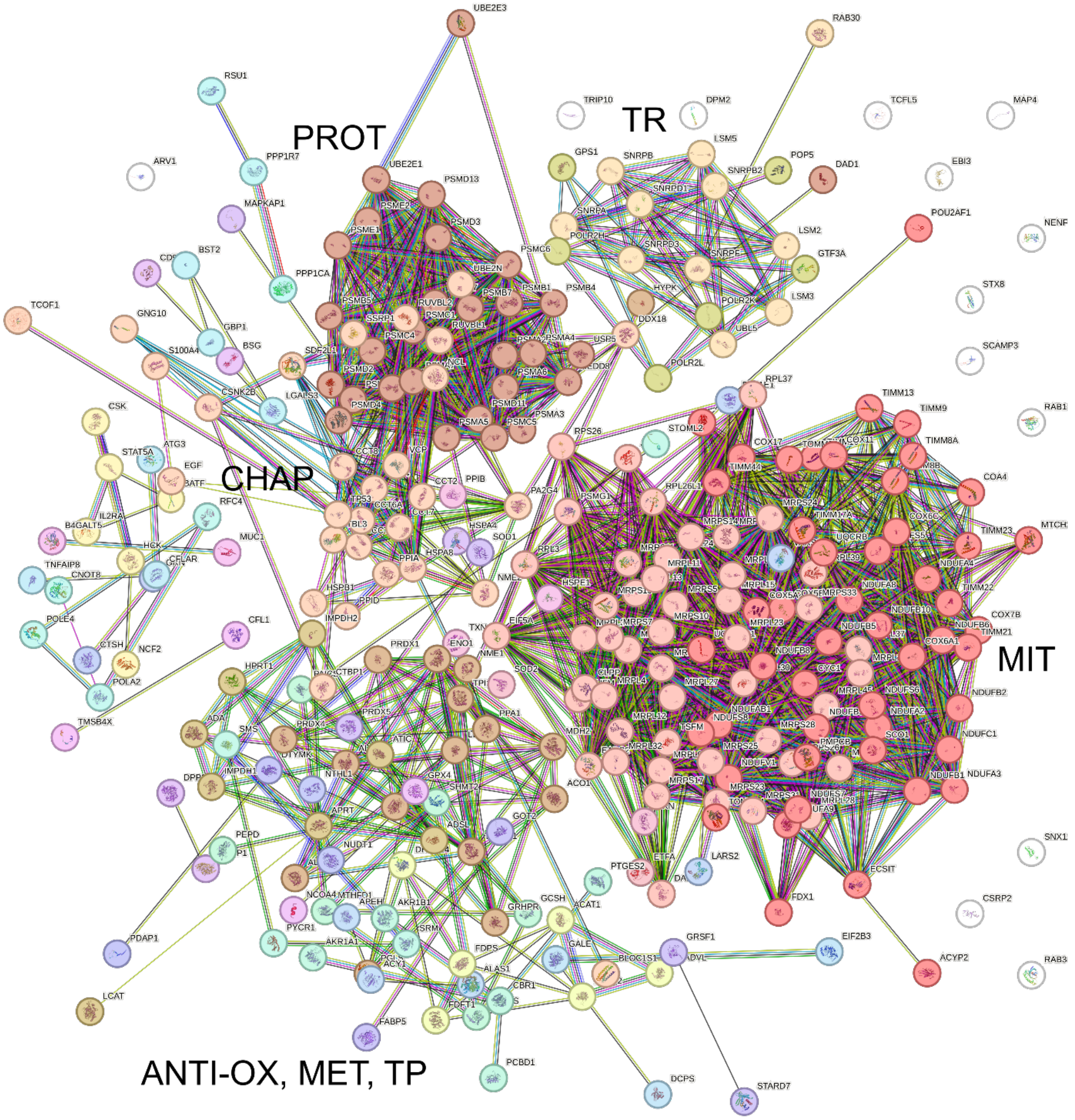


**Figure S2**. *STRING cluster analysis of genes upregulated after 22 h incubation of human CD19$^+$ B cells with IMT504*. Functional clusters include mitochondrial-related genes (MIT), transcription-related genes (TR), proteasome components (PROT), chaperone proteins (CHAP), antioxidant defense genes (ANTI-OX), metabolism-related genes (MET), and tissue protection genes (TP). Notably, ANTI-OX, MET, and TP clusters exhibit limited connectivity and appear interspersed, suggesting minimal interdependence and potential for independent regulation under cellular stress.
